# Functional significance of spectrotemporal response functions obtained using magnetoencephalography

**DOI:** 10.1101/168997

**Authors:** Francisco Cervantes Constantino, Marisel Villafañe-Delgado, Elizabeth Camenga, Katya Dombrowski, Benjamin Walsh, Jonathan Z. Simon

## Abstract

The spectrotemporal response function (STRF) model of neural encoding quantitatively associates dynamic auditory neural (output) responses to a spectrogram-like representation of a dynamic (input) stimulus. STRFs were experimentally obtained via whole-head human cortical responses to dynamic auditory stimuli using magnetoencephalography (MEG). The stimuli employed consisted of unpredictable pure tones presented at a range of rates. The predictive power of the estimated STRFs was found to be comparable to those obtained from the cortical single and multiunit activity literature. The STRFs were also qualitatively consistent with those obtained from electrophysiological studies in animal models; in particular their local-field-potential-generated spectral distributions and multiunit-activity-generated temporal distributions. Comparison of these MEG STRFs with others obtained using natural speech and music stimuli reveal a general structure consistent with common baseline auditory processing, including evidence for a transition in low-level neural representations of natural speech by 100 ms, when an appropriately chosen stimulus representation was used. It is also demonstrated that MEG-based STRFs contain information similar to that obtained using classic auditory evoked potential based approaches, but with extended applications to long-duration, non-repeated stimuli.

**Author summary:** The spectrotemporal response function (STRF) model of linking dynamic acoustic stimuli to dynamic neural responses is applied to whole-head non-invasive magnetoencephalography (MEG) recordings of the human auditory cortex. MEG STRFs were consistent predictors of neural activity, quantitatively and qualitatively, by comparison to those obtained from animal models using local field potential or multiunit activity as neural responses. Comparison of STRFs using stimuli as diverse as tone clouds, natural speech, and music revealed a common structure consistent with shared baseline auditory processing, when an appropriately chosen stimulus representation was used.

## Introduction

Empirically measured sensory receptive fields and response functions offer analytical characterizations of computations attainable by the auditory system[1–3]. Applied linear systems methods such as the spectrotemporal response function (STRF)[4–6] have similarly led to informative computational characterizations of central auditory neural function with respect to sound encoding and perception[7]. The STRF can be viewed as a representation of the approximate neural response to changing auditory input in time or frequency; any particular functional description will vary according to the location and role of the neurons. Different stimulus classes (e.g. artificially generated sounds vs. natural sounds) may produce related, but dissimilar STRFs from the same neural unit, speaking to fundamental processing differences (and similarities) of auditory encoding[8–10]. An emerging view in electrophysiology is that the STRF may represent a snapshot of the entire network converging onto that neuron (or group of neurons)[10], incorporating this population’s activity in its neural representation of the spectrotemporal features of the stimulus[7]. As seen here, STRFs also have a role in investigations of ensemble auditory coding, using neural recordings obtained from magnetoencephalography (MEG) or electroencephalography (EEG).

STRFs directly characterize the relationship between a sound stimulus and the accompanying neural response. For neural ensembles, rather than individual neurons, many individual linear components may be jointly pooled, perhaps even superadditively (depending on the underlying neuroanatomy and neurophysiology of the signal source). Also, as in the case of a single-neuron-based STRF, it may be methodologically simpler to use controlled stimuli rather than natural sounds[11–13]. It remains to be determined how the spectrotemporal features of ensemble-based STRFs correspond to the time-varying evoked-related-potential responses (and other standard MEG/EEG measures) as a function of frequency, and also to what extent the STRF encoding model can provide analogous additional information besides predictive power. Furthermore, the STRF estimate of a stimulus-response relationship may depend on the particular representation chosen for that stimulus; in particular, it remains unknown which specific stimulus representations are optimal for the purpose of matching STRF features to neural function[14], and whether any such choices can address the key question of how to generalize across stimulus classes, from artificial to natural stimuli. Finally, it is important to discuss overlap between these non-invasively obtained STRFs and those available from local field potential (LFP) data or from other invasive recordings.

In order to address these questions, evoked cortical activity recordings from healthy listeners were obtained with MEG during active listening of pseudo-random multi-tone patterns[12,15] presented at a variety of rates. STRFs were obtained per subject and condition, in order to assess the extent to which the MEG responses were linearly explainable by a sparse representation of the stimulus sound pattern, and whether rate-related changes are consistent with those found using invasive electrophysiological techniques. Peak components in STRFs and temporal response functions (TRFs) were identified and their latencies compared to those obtained with standard tone-based averaging. Alternative representations of the stimulus, including the auditory spectrogram, were used for reverse correlation in order to constrain the space of stimulus representations given the properties of the MEG cortical signal. Finally, these functionally informative STRFs were compared to those from datasets from studies using natural speech[16] and music processing[17]. This allowed an investigation of ensemble-dependent issues arising from STRF comparisons when using artificial vs. natural stimuli[9].

MEG-based STRFs are shown to functionally explain considerable amounts of response variability while revealing a parsimonious mapping of response features seen in classic averaging methods to those obtained from dynamic stimuli timeseries. Quantitatively, the MEG-based STRFs account for similar levels of predictive power to single and multiunit responses in auditory cortex[11]. Qualitatively, the mappings show reasonable correspondence with those from local field potential activity in animal models[18,19] and manifest similar stimulus dependencies (e.g., density[12]). We find that similar STRF structure is seen across responses to stimuli as diverse as natural speech and music, demonstrating convergence across stimulus classes. This last result, however, depends on the use of a specific (sparse) representation of acoustic stimuli, the nature of which provides additional knowledge regarding the role of spectrotemporal modulations on predictive frameworks of auditory cortical representations over a wide range of dynamic sound classes.

## Results

### MEG cortical responses predictable from the STRF linear model

Potential successes of the STRF as a linear model to predict MEG responses from acoustic stimuli are evaluated by comparing the actual vs. predicted responses which, unlike spike-generated STRFs, are continuous waveforms (Fig 1A). Model predictions are obtained by linearly convolving the corresponding STRF with the stimulus representation, using cross-validation (arbitrary separation of training data from testing data) to prevent overfitting, which makes this a conservative estimate due to noise present in the testing data[11,13]. If instead only the training data is used, i.e., fitting to the same data as is tested, STRF estimates provide a stringent upper limit as to how good any linear prediction can be. STRFs estimated using cross-validation predict the large negative deflections (Fig 1a, red) that follow tone onsets (∼100 ms post impulse) well, but unlike those from training (Fig 1a, blue), are less accurate for positive excursions (both data sets summarized in Fig 1b). The ability of the STRF model to predict the encoding relationship between sound patterns and cortical responses can be measured as the fraction of response variability explainable by the linear model, estimated on an individual condition and subject basis, once intrinsic response variability (unrelated to the stimulus) has been removed[11,13]. MEG STRF predictions were found to range as high as 34% of variance explained across participants and rates, using cross-validated data. When the fraction of variance explainable by the model was compared with normalized noise power (or inverse SNR), the explainable fraction in the theoretical noiseless limit was estimated to be 23.0 ± 2.0% (mean ± st. dev.; CI: 19.0–26.9%) as part of a significant linear regression relationship (*F*=45.9; *p*=2.7×10^−9^; *R^2^*=0.386), with an upper limit of 71% as provided by training-data only results (Fig 1c).

**Fig 1.**
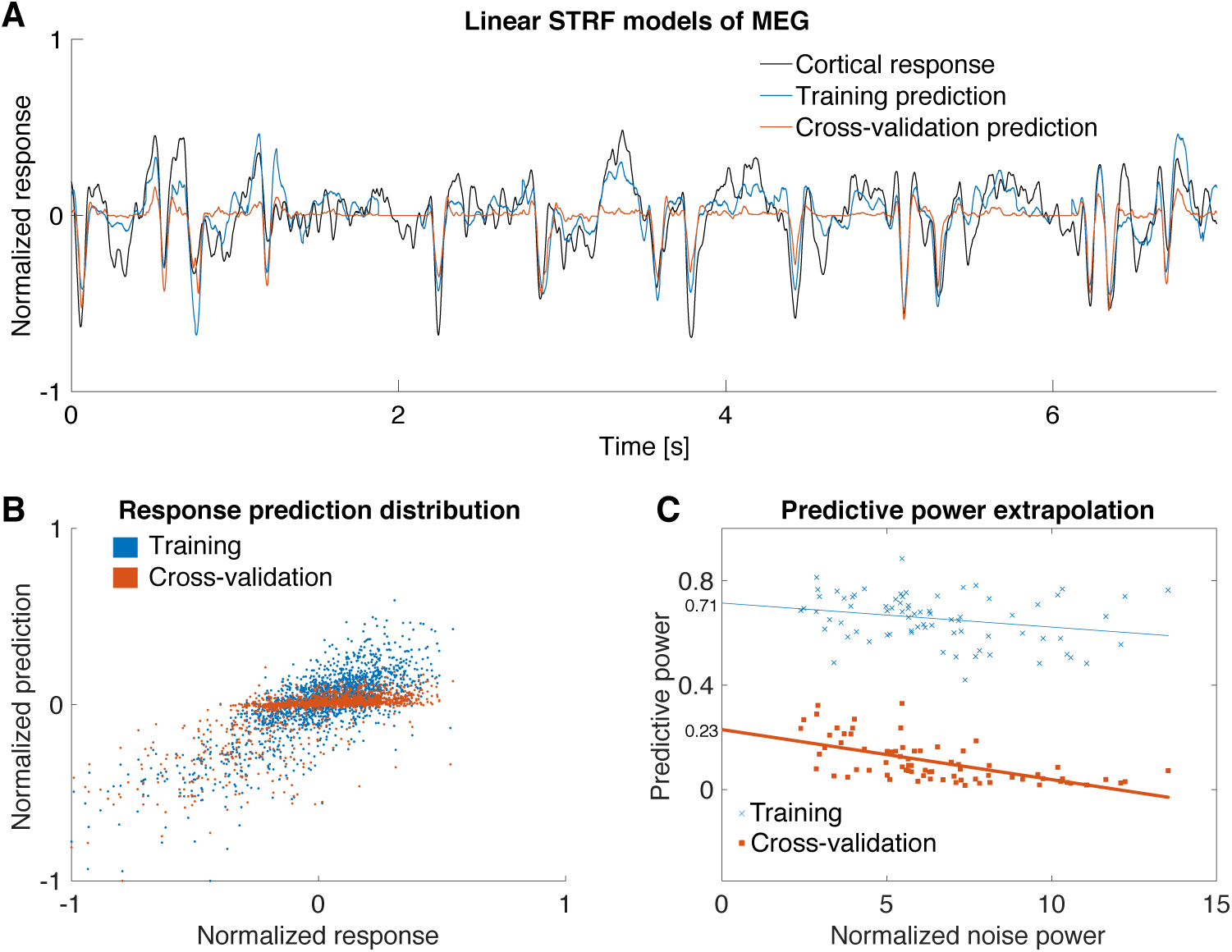
Spectrotemporal encoding models of MEG signals from human auditory cortex. a) A 7 s sample recording of an MEG response to a sparse multitone pattern (2 tones/s), with STRF-based predictions. b) STRFs were optimized by iteratively minimizing prediction error on the entire dataset, referred to as *training* (blue, *r*=0.74), or alternatively on their ability to generalize (*cross-validate*) over testing datasets (red, *r*=0.62). c) The predictive power of the STRF models is shown by linear regression of individual STRFs across participants and conditions on their corrected normalized noise power (i.e., inverse signal-to-noise ratio, an indicator of trial by trial reliability, see *Methods*). Extrapolation of performance to zero noise power gives a noise-corrected expected performance for both the conservative cross-validation-based estimates and the fundamental-upper-limit training estimates.

### Fraction of response explained by STRF features consistent with standard evoked potentials

STRFs based on MEG responses display consistent spectrotemporal structure in the form of positive-negative-positive complex deflections (Fig 2a) coinciding with typical auditory cortical latencies (e.g. those of the P1-N1-P2 complex in averaged EEG responses to isolated tones). In particular, the multitone STRFs demonstrate strong negative responses at ∼100 ms post impulse onset (STRF_100_). The specific STRF_100_ latency depends on stimulus frequency, varying ∼20 ms over the frequency range 180-700 Hz; at higher frequencies the latency is approximately constant (STRF_100_ latencies for 2 tones/s shown in Fig 2b, black). STRF_100_ latencies were found to follow standard tone-evoked M100 latencies[20–25] obtained under various conditions (Fig 2b; also Table 1). The correspondence suggests a quantitative link between the STRF_100_ and M100, and therefore between STRF-based techniques and ordinary auditory evoked cortical potentials. Analyzing the same experimental data using standard evoked response analysis instead (epoching and averaging over responses to all tones in the sparsest multitone pattern) demonstrates strong temporal correspondence at the group level (S1 Fig).

**Fig 2.**
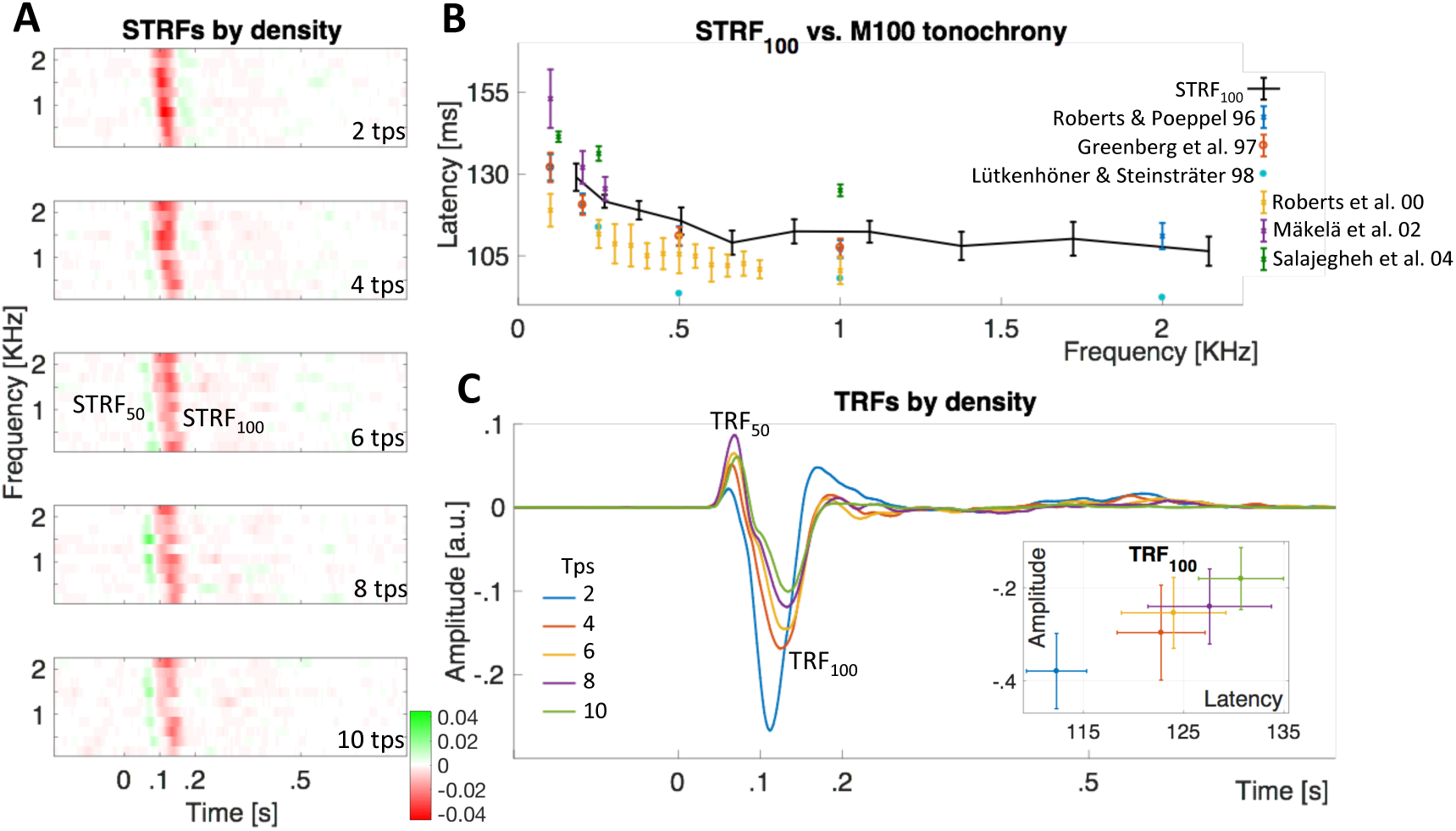
Consistency between response function model predictive model features and evoked potentials. a) Grand average spectrotemporal response functions based on multitone stimuli demonstrate a positive-negative-positive structured sequence between 50 and 200 ms following tone onset; tone cloud density introduces qualitative changes in relative amplitude and delays: at increased rates an early positive component (50-100 ms; STRF_50_) emerges, while the medium latency negative component (100-150 ms; STRF_100_) attenuates, and a late positive component (150-200 ms) present only in the sparsest conditions disappears. b) STRF_100_ components are delayed by over 20 ms as tone carrier frequency decreases from 2 to 0.2 KHz, in a manner consistent with those of evoked potentials in single tone presentations[20,21,23–25] (Table 1), indicating a correspondence between impulse response functions obtained through reverse correlation and averaged evoked potentials. A common latency decrease across studies and conditions is observed for carrier frequencies in the 180-700 Hz range. c) Temporal response functions, obtained by reverse correlation with the stimulus envelope collapsed across frequencies, show features similar to the P1-N1-P2 complex commonly found in EEG evoked potentials[26]. Higher tone presentation rates result in the emergence of the TRF_50_ and in decreased amplitude and increased latency of the TRF_100_ (inset), as well as the attenuation of a later-latency positive deflection. Error bars are 1 standard error of the mean.

**Table 1.** Comparison of studies reporting M100 absolute latency values in response to pure tones, with participants, recording mode, stimulus details, and M100 peak determination method where available.

|  | # of subjects<br>[mean age] | Sound delivery<br>[Sensor location] | Tone duration<br>(ms) | Presentation<br>rate [tones/s] | Peak finding<br>method |
| --- | --- | --- | --- | --- | --- |
| **Roberts & Poeppel, 1996[20]; Greenberg et al., 1997[21]** | 5<br>[24-33 y] | Monaural<br>[Contralateral, Left] | 400 | 0.7 - 1.3 | Equivalent dipole, Maximum RMS |
| **Lütkenhöner & Steinsträter, 1998[22]** | 1<br>[28 y] | Monaural<br>[Contralateral, Left] | 520 | ~0.4 | Maximum RMS |
| **Roberts et al., 2000[23]** | 8 [-] | -<br>[Both hemispheres] | - | - | - |
| **Mäkelä et al., 2002[24]** | 11<br>[32 y] | Binaural<br>[Both hemispheres] | 200 | 1 | Optimal sensor pair |
| **Salajegheh et al., 2004[25]** | 11<br>[45.8 y] | Binaural<br>[Both hemispheres] | 400 | 0.8 - 1.3 | Maximum RMS from optimal 12 sensors |

To further investigate the correspondence between STRFs and evoked potentials (specifically the effects of tone density), reverse correlation was performed with respect to frequency-collapsed representations of the stimulus, generating the frequency-independent Temporal Response Function (TRF, Fig 2c). The ∼100 ms latency negative peak (TRF_100_) amplitude decreased with increasing tone-density by ∼60% across modulation rate range studied, while latency increased 20% (see inset). In contrast, the ∼50 ms latency positive deflections (TRF_50_) had the smallest amplitude for the sparsest multitone condition. Thus sources with ∼50 ms latency generate a strong increase in cortical activity with a transition from scattered to continuous pure tones, while sources with ∼100 ms latency decrease in strength as they are delayed. Cortical activity in sources with 150 ms latency may also be active, provided the inter-tone interval is long.

### STRF most informative for onset-based representations of multitone stimulus

Methodologically, the acoustic representation of the stimulus used to generate the STRF may employ any number of available time-frequency representations of the sound, including the widely-used spectrogram[11,27–29]. One reason to consider alternatives to the spectrogram is to compare STRF features with evoked response features, since an evoked response to tones is calculated not with respect to the spectrotemporal duration of the tones but only to their onsets. Thus analyses also included binary and sparse representations of the stimulus: single tones were modeled as trigger-like impulses timed with tone onset and organized by frequency. Indeed, stimulus features that are known to be encoded by auditory cortex include onsets, offsets, and stimulus duration (in the form of sustained responses)[30–33]. Since the MEG signal is aggregated across synchronized individual neurons[34], evidence for those same encodings requires investigation. Reverse correlation techniques are well suited for this larger-scale analysis because it explores the outcome of alternative stimulus representations that emphasize such features. The stimulus representations tested here (cf. Fig 3 insets) were (i) the ideal *trigger* representation, (ii) the ideal *edge* representation (both onset and offset triggers), (iii) the ideal stimulus first-order *derivative* (onset and negatively-signed-offset triggers), which can itself be used to generate the trigger representation if followed by half-wave rectification, (iv) the ideal stimulus *pulse* envelope, which has constant value from onset to offset (and which can itself be used to generate the previous representation if followed by differentiation), (v) the actual acoustic stimulus passed through a filterbank with identical center frequencies as the tone, whose *envelope* is then extracted (see *Methods*), and (vi) a generalized *envelope onset* representation obtained via half-wave rectification of the previously defined filterbank envelope output. Only the last two can be applied to natural (non-discrete) stimuli, and so are especially important in later sections.

**Fig 3.**
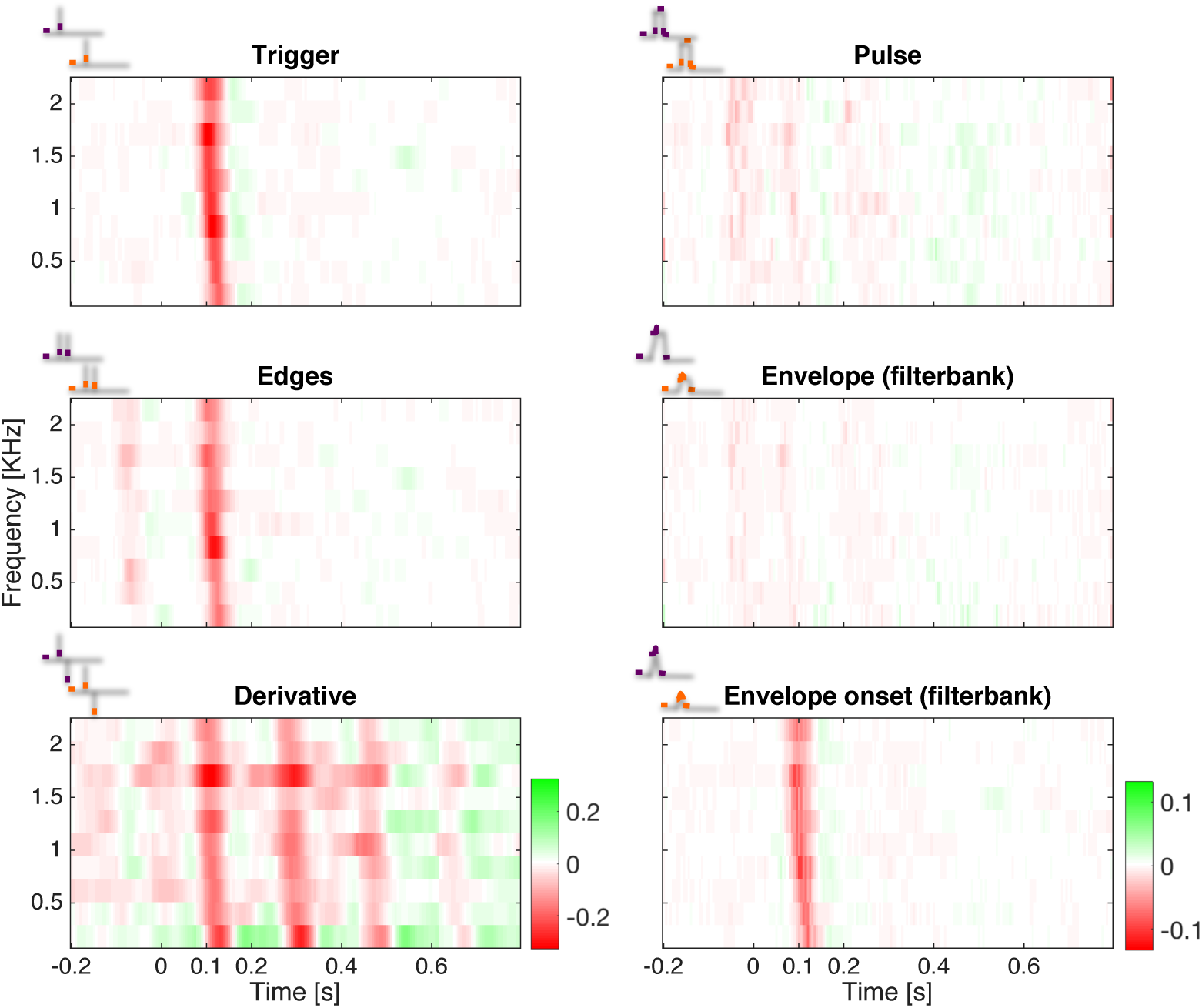
STRFs generated using different stimulus representations achieve different levels of functionality. STRFs generated from multitone patterns are functionally informative (e.g., comparable to evoked potential analysis) when each individual tone is discretely represented by its onset (top left) but not when represented instead by the timing of its temporal edges (middle left), sign (bottom left), or a discrete representation of the entire pulse duration (top right). Related to the spectrogram, the representation based on passing the acoustic signal through a series of filterbanks, then extracting envelopes per band (middle right) yields only barely discernible results. Extracting onset timing information from the filterbank, in contrast, was quite functionally informative (half-wave rectification of the first derivative of the filterbank output; bottom left). Critically, filterbank-based methods do not require *a priori* definitions of temporal edges and can be used for arbitrary stimuli. Color scales as in bottom right inset, except for Derivative STRF.

Grand average STRFs in Fig 3 demonstrate that among such representations, only those expressing tone onset events explicitly yield components comparable to those of evoked potential analysis (first and last STRFs of Fig 3); STRFs from the alternatives introduced ringing and/or pre-causal artifacts. As with the original onset-based trigger representation, reverse correlation with both temporal edges predicted activity from the first edge in accordance with the latency by frequency dependence, but also produced a pre-causal mirror component, in advance of the original and at the tone duration distance. This pattern suggests that tone offset was not explicitly encoded here. This interpretation is supported by analysis of STRFs generated by the derivative representation, which correspondingly flips the sign of the same pre-causal mirror component, but is additionally contaminated via constructive interference by a series of artefactual ringing cycles. The pulse representation, which can be viewed as an idealized envelope, produces STRFs that are essentially featureless (or at best, whose features are barely discernible above the noise floor). This result is unexpected since typical auditory reverse correlation studies use a duration-based stimulus envelope representation[35,36] and the temporal envelope is often hypothesized to be the response-driving feature. Similarly, the acoustic envelope representation (using a filterbank model; see *Methods*) also produced featureless STRFs. An attempt to re-create the onset representation (i.e. half-wave rectification of the acoustic envelope representation derivative), did however generate STRFs with features comparable to evoked potential analysis, and enables the extraction of onset-like information in general from diverse complex natural stimuli. Because of the remarkable agreement between the idealized and the acoustic onset models, interpretations based on evoked potentials may extend to reverse correlation analysis applied to other stimulus classes where definitions of onsets would be *a priori* unknown or not controlled for, such as natural sounds.

### Convergent STRF models across artificial and natural stimuli

Because of its potential to reveal hierarchical processing mechanisms, a major goal in auditory reverse correlation has been to examine the encoding relationship for critical natural stimuli including speech and other communication sounds. To this end, datasets from two previously unpublished studies on speech and music processing were submitted to the same analysis methods as the multitone pattern (Fig 4a), with stimuli represented by their envelope onsets. As with the onset-based representation of the multitone patterns, STRFs for speech and music exhibited qualitatively similar structures, with distinctive biphasic components near 50 and 100 ms post rising transient impulses (onsets) along the same investigated spectral region. Inspection of the stimuli under either envelope or envelope onset representations suggests that the latter procedure effectively increases similarity in the underlying distribution across stimulus classes (S2a Fig). The frequency dependence of relative peak delays was also maintained for these stimulus classes (S2b Fig) but with class-dependent timing differences, suggesting a common fundamental mode of spectrotemporal cortical processing up to ∼200 ms and after which notable processing differences appear according to the stimulus class. While neural data from all studies were obtained from different subject groups, one subject did participate in those two studies and in a modified pilot version of this experiment (2.4 tones per second presentation rate); these data are presented in Fig 4b, again showing strong qualitative similarity both to group data and class-dependent timing differences. This subject’s topographic magnetic field maps associated with the neuromagnetic signals derived in each of the three studies are displayed in Fig 4c; mapping each STRF to overlapping spatial distributions is consistent with source activity at the superior aspect of the temporal lobes.

**Fig 4.**
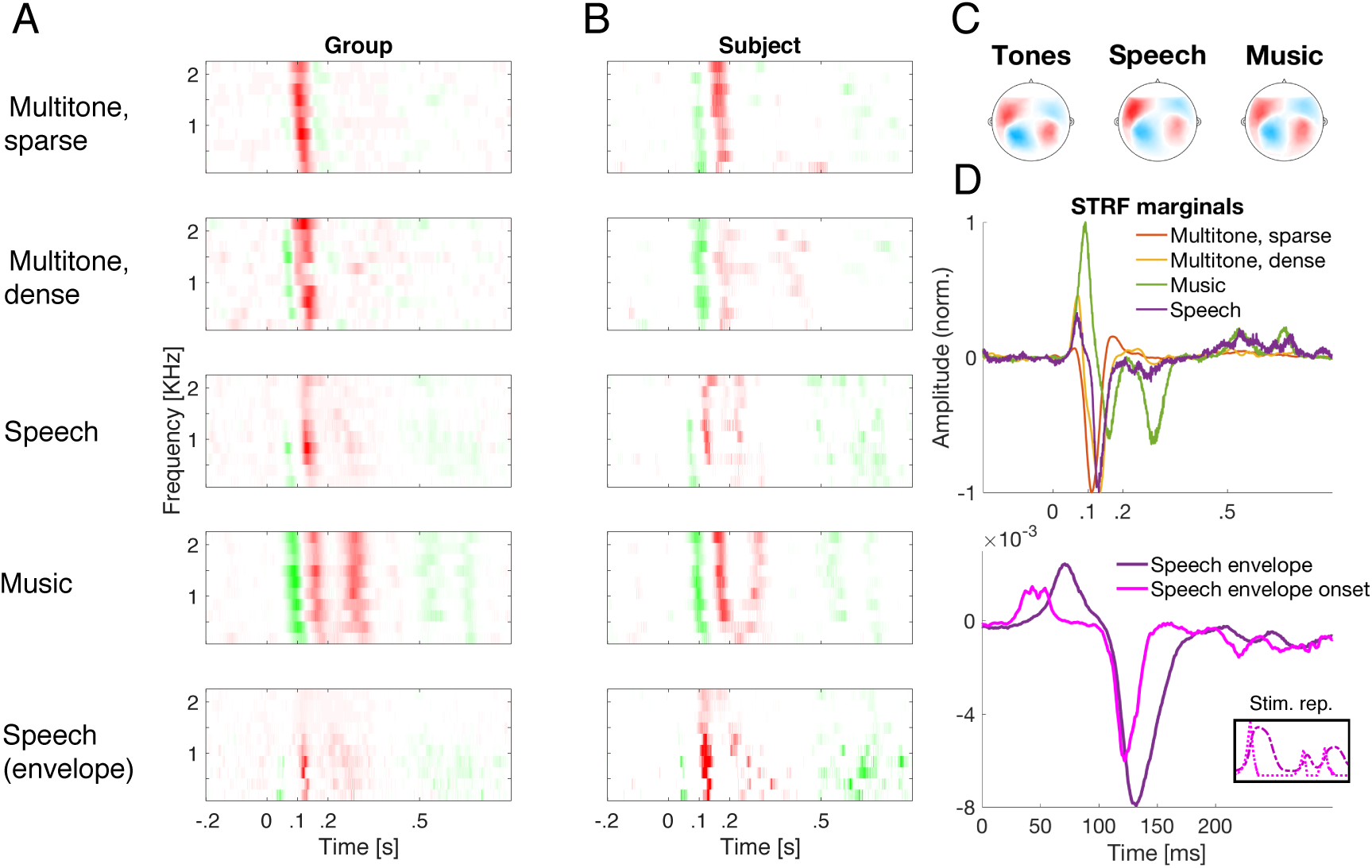
Interpretational power from stimulus representations across STRFs from different stimulus classes. a) Group normalized STRFs from the multitone pattern experiment (*N*=15), and from studies on natural speech (*N*=12) and on music (*N*=15), reveal considerable structural similarity when stimulus onset is extracted as a driving feature of the neuromagnetic response. b) Neuromagnetic STRFs from the same participant across the tones, speech, and music studies, which show substantial consistencies across stimuli when represented by their temporal envelope onsets per frequency band. c) The topographic distribution from same subject as in (b) revealed strong bilateral consistency across classes but with increased left hemisphere-bias during speech processing. d) Top: Timing of major neuromagnetic activity peaks, as shown by TRFs derived from spectral integration of the STRFs in (a), results vary depending on stimulus class and/or context: earliest positive and negative deflections change with increasing acoustic density but also with additional spectrotemporal complexity as found in natural speech and music. Bottom: Group TRFs comparing both speech envelope and envelope onset related activity. Timing differences are explainable by differential acoustic representation in early (< 0.1 s) but not late activity peaks, suggesting the formation of higher order neural representation of elements in speech acoustics by ∼100 ms. Only the first deflection timing difference is explained by slope-to-maximum time differences between stimulus representations (inset, same color coding). Curves smoothed by a 5 ms moving average.

To better illustrate class-dependent temporal differences across the studies, TRFs were obtained by collapsing STRFs across spectral bins, as shown in Fig 4d. These plots emphasize spectrally consistent changes in temporal processing due to stimulus class, along with relative amplitude differences. As before, early activity appeared least prominent for the spectrotemporally sparsest stimuli; in the case of the single participant tested across all three stimulus classes, a high-temporal resolution analysis of the multitone TRF_100_ shows its dynamics are very close to those of the speech envelope counterpart (S3a Fig), with characteristic time constants of ∼3 ms (S3b Fig). The response dynamics for music, however, do not follow similarly, which suggests that features other than overall acoustic onsets may contribute to synchronized auditory responses in these cortical populations.

### Cortical transformation of natural speech envelope representation

In reverse correlation analyses, exploration of alternative representations of the stimulus may provide complementary insight into the functional operations by the auditory system. Fig 4a and Fig 4b show that for natural speech, STRFs based on the acoustic envelope (row 5) led to functionally informative STRFs, consistent with prior approaches[36,37]. STRFs based on the envelope *onset* representation (row 3) are similar, which is expected since the envelope onset is correlated with the original envelope. In terms of timing, the corresponding group TRFs (Fig 4d) show a difference of 43 ms between TRF_50_ peak components. This was found to be the same as the characteristic delay between their underlying representations, obtained by cross-correlation of the stimulus representations. Such a close correspondence is evidence that at the level of the neural source of the TRF_50_, an increasing acoustic envelope operates as a fundamental auditory feature of the stimulus. In contrast, the corresponding comparison of STRF_100_ peaks across the two representations (envelope and envelope onset) gives a much reduced difference of 20 ms (Fig 4d, S5 Fig), not consistent with the acoustic differences between the corresponding representation peaks. Compression in components’ relative delays were observed across spectral bins (S2b Fig), as well as in individual temporal response functions (S3a Fig, S4 Fig).

## Discussion

The present investigation describes STRFs as a series of response function mappings from artificial and natural sounds to auditory neural responses. It has been demonstrated that these STRFs possess similar predictive power as their single-unit cortical counterparts, and, importantly, show strong similarities across stimulus classes when an acoustic envelope onset representation is chosen. Specific choices of spectrotemporal stimulus representations[14] result in STRF models that are not only predictive but whose temporal structure is highly consistent with that from standard evoked potential components.

### Comparison to spike-based spectrotemporal receptive fields

The spectrotemporal *receptive field* can be considered a spike-triggered averaged spectrogram, from auditory periphery[6,38] or central nervous system recordings[2,39–41]. Since reverse correlation is a more general principle than spike-triggered averaging[42] it has been used here to characterize and predict the neural responses of auditory systems where both input and output are continuous time-series[43] via the underlying *response function* of the system. Whether measured by spikes or continuous neural responses, neural systems are non-linear, so predictive linear models of central neural coding are necessarily incomplete descriptions of the underlying coding relationship and are bounded by the predictive power and interpretability they maintain within the limits of the linear regime[11].

The multitone stimulus employed here is comparable to a dynamic random chord stimulus[11,44,45] though it has more temporal degrees of freedom, allowing cross-frequency overlap in a continuing pattern that prevents constant tone presentation rates. It is more similar to dynamic random chords than other artificial stimuli used to estimate STRFs, such as ripple noise and moving ripples[46,47], which focus on stimulus modulations instead. The predictable fraction of variance in the evoked MEG source timeseries was found to be 19–27%, in close correspondence with 18%[11] and 31%[48] predictive power from primary auditory cortex (A1) single/multiunit responses. Comparisons regarding predictive power (and other STRF properties) should also take into account fundamental differences in the underlying signal (spiking versus dendritic-origin activity) and its scale (neuron or highly local population versus meso-scale cortical patches[34,49]), the animal model, and state (e.g., performing a task vs. resting vs. anesthetized)[7].

Qualitatively, the STRFs presented here exhibit a general broadband structure with frequency-dependent latencies and amplitude changes depending on stimulus density. Remarkably, similar properties appear in STRFs obtained from LFP in mammalian A1[18,19,50], featuring broadband inhibitory-excitatory component sequences and, often, frequency-dependent latencies. Component latencies in mammalian A1 are ∼50% shorter than here, which may be explained by reduced equivalent cortico-cortical transmission length delays[51] for the species involved in those studies. With respect to human studies, the component latencies reported here are consistent with multiunit activity[52] and high-gamma activity in electrocorticography (ECoG)[45] in the functional equivalent of A1, Heschl’s gyrus. The STRFs obtained in such datasets principally reflect neural spiking, resulting in mappings with narrow-band features, consistent with their interpretation as units locally sampled along the tonotopic gradient of A1. Indeed, frequency selectivity becomes reduced for local field potential recordings[18,50] (i.e. ECoG frequencies below high-gamma) as they sample redundant activity across distant recording sites with intra-cortical interactions[53] – which may effectively smooth the spectral selectivity distribution[18]. Unlike local recordings, which due to high-frequency selectivity can require that receptive fields be realigned by best frequency[28] to extract statistical features, MEG STRFs offer distributed access to more global cortical network domains. Plausibly, analog results may be expected from future human auditory LFP STRF studies from invasive procedures, given that these have typically focused on multiunit and high-gamma activity[45].

In addition, these MEG-based tone-generated STRFs show stimulus-dependent differences as seen elsewhere in the receptive field literature (see review by Eggermont[7]), namely, amplitude decreases with density. This is consistent with awake primate results, where three-fold increases in tone presentation rate (9.7 to 31 tones/s) may be accompanied by a magnitude decrease of about a third in the STRF maxima[12]; here, a similar multiplicative change in tone density (2 to 6 tones/s) produced a peak decrease of about half. Suppression of excitatory contributions[15], or emergent inhibitory activity throughout A1 single units[12] have been proposed as mechanisms for response field modulations observable from LFP recordings[50]. In cortical neurons, increased firing rates may accentuate depression rate imbalances between excitatory synapses and those with increasingly inhibitory activity[12,19,28], which is a known factor involved in receptive field modulations in somatosensory[54] and visual areas[55]. For auditory recordings, more inhibition may effectively increment responses’ spectral specificity or bandwidth at higher tone densities[12,50] – the analog of which was not observed in the MEG STRFs (see Westö & May[56] for a cautionary note on interpreting inhibitory contributions to STRFs following dense stimuli). Among factors reducing STRF predictive power is the increase of inhibitory fields in estimates from single unit recordings[12]; in MEG this effect appeared to be mirrored by response function components of opposite sign to the STRF_100_. Further research is thus necessary concerning the coarse-grained level of analysis that is accessible via MEG/EEG respectively, in comparison to that afforded by single/multiunit signals.

### Association with auditory evoked potentials

Unlike traditional averaging methods, reverse correlation involves continuous delivery of a dynamic stimulus in order to generate a predictive model (of novel instances of the same sound class). It has been shown here that STRFs and TRFs can be directly compared to standard auditory evoked responses, namely the magnetic M50, M100, late auditory evoked responses, and the P1-N1-P2 complex in EEG[26]:

(i) The earliest positive-polarity component, the STRF_50_, seen at higher multitone densities, is a temporal analog to the M50 response originating from Heschl’s gyrus (including core/primary areas)[57,58]; its amplitude may also be modulated by interstimulus intervals[58] at low presentation rates (<2 tones/s). Known modulators of M50 amplitudes include harmonic versus noise-like bursts[59,60], prepulse inhibition[61], and automatic processing of redundant information as a form of sensory gating in paired-click stimulus designs[62,63]. In terms of predictive power, this component did not generalize well over novel instances of the multitone random pattern; this is consistent with an adaptive role contributing to considerable changes in the response profile dependent on the local context on the order of a few seconds or less.

(ii) The subsequent major component, the negative-polarity STRF_100_, exhibited magnitude decrease and delay increase with density, with a sharp transition after the sparsest density level. Suppression of the M100 response from supratemporal cortex has been observed in the transition from low to higher tone presentation rates[58] highlighting the interpretation of increased inhibitory effects that include generalized refractoriness among neurons at denser conditions[64]. This component is also subject to attentional modulation[65–67] which may reflect that individual tones in a densely populated scene fail to capture attention individually. Because of this component’s involvement in tracking perceptual objects of an auditory scene[37], and of the increasing quality of flow and continuity in these artificial stimuli, sharp transitions in this component may suggest indices of ‘crowding’ relevant to the figure-ground separation problem[68,69]. Accounts of spectrally-dependent latency in the evoked M100 components[20–25] were consistent at this stage and fall within the sensitivity domain for human voice pitch production and discrimination[24]. These latencies are also consistent with those of pitch-specific onset responses, whether elicited by complex tones or by centrally-generated Huggins pitch percepts[70,71].

(iii) The second major positive-polarity peak appears only in response to the sparsest stimulus. Auditory event-related potentials at ∼200 ms latency have been described in EEG as expectancy indices, exhibiting greater amplitudes for tones whose presentation in time is uncertain[72]. At denser conditions, shorter inter-stimulus-intervals may reduce the tone-evoked analogous EEG P2 component, regardless of presentation within a repetition sequence or as an oddball, suggesting involvement of modulation mechanisms other than habituation[64].

### On alternative representations of stimulus state

In addition to their predictive power, STRF profiles are functionally informative in a way similar to trial-averaged evoked responses to isolated stimuli[8,14,73]; this was the case for STRFs compared across stimulus classes when the stimulus representations were filterbank-derived onsets. Other abstract representations of this stimulus pattern, including both temporal edges, their directionality, the duration of sustained acoustic energy, or, related to the latter, the spectrogram, did not appear to be similarly functionally informative – even though they contained and extended information from onset representation. Nevertheless, predictive power was similar across representations, suggesting that this metric alone is insufficient to expose which aspects of the stimulus map to the system’s response. More complex tone patterns might allow predictive power to become more informative regarding the statistical characterization of a stimulus (c.f.[74]). The lack of evidence for explicit neural encoding of offsets is in accord with neurophysiological evidence suggesting offset-encoding cells to be outnumbered by onset cells, and/or to have minor neural response profiles relative to onset encoders[30,31], which in the aggregate would result in differential contributions to the neuromagnetic response.

### On extension to natural stimuli

Processing of environmental sounds, including conspecific calls, is a critical auditory task. Encoding models incorporating natural sounds with complex spectrotemporal structure provide powerful computational insights into the auditory system that may be inaccessible with synthetic stimuli only[7–9]. STRFs derived from invasive recordings from A1 perform similarly in terms of predictive power, using random tone chord stimuli, animal calls, environmental sounds, sound effects, and music[11,27], at the population level. For some subset of these neurons, successful linear encoding of the spectrogram may also occur in the same unit for *both* artificial *and* natural vocalization encodings[10,28]. The search for predictive models that generalize over novel stimuli not in the training set has proven difficult however[10,44]. The temporal statistics intrinsic to natural sounds may be critical[10], and some evidence from A1 STRFs demonstrates higher predictive power using conspecific vocalizations that are not dilated or compressed[75]; similarly, comparisons are also favorable for artificial and communication sounds controlled for the span of their temporal and spectral modulations jointly, but allowing differences in their amplitude fluctuations over time[10]. Observed stimulus-class dependencies in STRF spectrotemporal properties appear as small time-shifts of STRF features, plus the emergence of additional late activity for speech and music. Analysis of such differences suffer from confounds arising from statistical nonuniformities among the sampled classes[8–10,29] and fully addressing this issue is beyond the scope of this investigation. The question of whether detailed class-dependent temporal coding frameworks may be achieved by means of linear methods remains open.

### On speech-derived STRFs

Advances in understanding cognitive processes relevant to speech processing have followed from reverse correlation studies that used the speech acoustic envelope (as represented by low-frequency, 1-15 Hz fluctuations in ECoG[76,77] and MEG/EEG[36,37,78,79] recordings). We find that the speech envelope STRF_100_ component exhibits similar spectral-dependent latency as the M100 evoked response, thus suggesting a level of speech analysis that still contains independent spectral information. Although in contrast with findings of near-constant M100 latencies for certain synthetic vowels presented in isolation[24], reverse correlation methods over long natural speech presentations are better suited to probe domain-general processing in realistic conditions due to their extended sampling,.

Additionally, the methods may constrain the time course of the change in neural representations of human speech from spectrogram to higher level. The low-frequency speech envelope and its onsets are both operationally related to functionally informative STRFs. The systematic delay between timeseries (peaks in the latter systematically precede those in the former) directly accounted for the relative difference between the resulting pair of STRF_50_ components after each representation. The acoustic mismatch could not explain, however, the reduced relative difference between subsequent STRF_100_ pairs. The interpretation of a compression is consistent with current models of step-wise speech processing, where the formation of speech analysis units or objects is preceded by an earlier spectrogram-like representation of acoustics completed by ∼80 ms post speech impulse onset. After this time, response functions did not account for the expected mismatch, suggesting a neurally-based progression into a modified stage of the neural representation of speech and adding to a body of MEG evidence for a cortical hierarchy of speech object representations (see review by Zhang and colleagues [80]).

Overall, these results demonstrate an important advantage of STRFs over standard epoch-averaging methods commonly when used in MEG applications, e.g., characterizing the phenomenology of disorders in clinical populations[81]: their ability to generalize to critical sounds beyond pure tones, most importantly natural speech. By providing both neural predictions and functional information, it allows noninvasive approaches to understanding developmental[82], learning and associative effects induced by tasks[83–85], or behavioral contexts[86,87] – thus potentially furthering insight into the role of dynamical representations of sound in auditory cognition.

## Methods

### Participants

15 subjects (6 women, 23.2 ± 2.9 years of age [mean ± SD]), 1 left-handed[88], participated in the multitone study. 12 subjects (6 women, 24.1 ± 3.0 years of age), all right-handed native English speakers, participated in the speech study. 15 subjects (5 women, 21.0 ± 1.7 years of age), all right-handed, participated in the music study. Each subject received monetary compensation proportional to the study duration (approximately 1.5 hours). Subjects had no history of neurological disorder or metal implants. The experimental protocol was approved by the UMCP Institutional Review Board and before each study session, informed written consent was obtained from the participant.

### Stimuli

#### Multitone study

Sound stimuli were constructed with the MATLAB® software package (MathWorks, Natick, United States) at a sampling rate of 44.1 KHz, and consisted of 50 s auditory scenes composed of pseudo-randomly presented 180 ms tones, each with frequency *f_i_* taken from a pool of 10 fixed values (range: 180-2144 Hz) in 2 equivalent rectangular bandwidth (ERB) steps[89] specified by *f*_*i*_ = *f*_*i*−1_ + (24.7 (1 + 4.37*f*_*i*−1_/ 1000)). For each frequency, tone onset times were uniformly distributed with a minimum inter-tone gap of 40 ms. Five tone presentation rates (2, 4, 6, 8, or 10 per second over all channels) were used separately. Tone onset times *T*_*j*_ were independent across frequency bands and selected in 20 ms bins. Individual tones were modulated with 10 ms raised cosine on- and off-ramps. Tone level was calibrated according to frequency based on the 60-phon normal equal-loudness-level contour (ISO 226:2003) in order to adjust for perceived relative loudness differences; relative gains to a 1 KHz reference were determined in 2 dB SPL steps. *Speech and music studies*. For the speech study, a 60 s female voice audiobook excerpt [90] narrated from *The Light Princess* (Macdonald, 1864) was used as part of a related study on reverberant speech processing [16]. For the music study, 55 s samples across 6 different instrumental musical styles reflecting a variety of genres and traditions, were presented: orchestra, *Symphony in F Major, No. 32, Movement I* (Sammartini, c. 1740); swing, *Cascades* (Combelle, c. 1940); blues, *Blues for B&W* (Rogers & Hilden, 2003); sarangi, *Raga Mishra Bhairavi: Alap* (Narayan, 2002); pipa, *Dance of the Yi People* (Huiran, c. 1960); and a euphonium transcription of *Dancing Night Wind* (Benning, 1997).

In all studies, audio signals were normalized and presented through the Presentation® software package (NeuroBehavioral Systems, Berkeley, United States), using audio equipment equalized to a transfer function approximately flat from 40 to 3000 Hz. Sound stimuli were transmitted to subjects via ear insertion tubes E-A-RTONE® 3A of 50 Ω impedance and E-A-RLINK® disposable foam intra-auricular ends (Etymotic Research, Elk Grove Village, United States) that were inserted in the ear canals.

### Experimental design

For the *multitone* study, trials consisted of a main tone cloud pattern scene presented in series with per block, generated anew per each subject. This resulted in trials that contained between 0 and 3 multitone density transitions within the trial, and ranged from 70 to 120 s duration. Each of the five main scenes were repeated 4 times, and only these data epochs were analyzed. After a brief training session, subjects were instructed to attend to the ongoing stimulus with their eyes closed and to report rate transitions via a button press. Optional rests were available every 5 trials, totaling 1.5 hours recording time. Subjects received feedback on the correct number of transitions at the end of each trial. For the *speech* study, trials consisted of various story passages presented in random order at different reverberant noise levels. At the end of a trial, subjects were asked comprehensive questions about the passage, and rated its intelligibility. For the present study purposes, analysis was based exclusively on reverberation-free, no-noise (‘clean’) trials, repeated 3 times across the experiment. For the *music* study, trials consisted of each of the 6 samples presented individually in random order. At the end of each trial, a 5 s clip taken from the same or a different piece was presented and subjects identified if it was an excerpt of the preceding trial. Each sample repeated 3 times across the experiment.

### Neural data recording

Magnetoencephalography (MEG) data were collected with a 160-channel system (Kanazawa Technology Institute, Kanazawa, Japan)[91] inside a magnetically-shielded room (Yokogawa Electric Corporation, Musashino, Japan) at a sampling rate of 1 KHz. Superconducting quantum interference device (SQUID) sensors (15.5 mm diameter each) were uniformly distributed (∼25 mm) inside a Dewar vase containing liquid-He refrigerant, with a concave outer surface fit to the average human head. Sensors are first-order axial gradiometers with 50 mm separation and sensitivity greater than 5 fT·Hz^−1/2^ in the white noise spectral region (> 1 KHz), except for three additional reference magnetometers separated from the neural sensors and arranged orthogonally to each other. A 1 Hz high-pass analog filter, a 200 Hz low-pass analog filter, and a 60 Hz analog notch filter were applied online respectively. Sensor channels with saturating or zero responses over more than 12.5 s recording time were excluded from analysis. Participants laid supine inside the magnetically shielded room and were asked to minimize body movement, particularly from the head.

### Neural data processing

#### Environmental noise

To eliminate environmental magnetic noise contributions, time-shifted principal component analysis[92] (TS-PCA) was applied, a process that discards optimally-filtered environmental signals recorded on the reference sensors. Reference sensors were 3 physical magnetometers (see *Neural Data Recording*) plus 2 virtual channels obtained by independent component analysis[93] of the remaining data sensors and selecting the two components with the most unstructured broadband (0-500 Hz) power. *Sensor-specific noise*. Electronic sensor noise was removed via sensor noise suppression (SNS)[94] by substituting each channel signal with its projection onto the orthogonal basis space generated by all other sensors in the system. This method exploits redundant activity across elements of the dense array (where the number of channels exceeds the number of brain sources of interest) by attenuating components specific to any single channel. *Spatial filtering*. Data-driven spatial filters were derived per participant using responses evoked by repeated trials in each of the respective studies. Response epochs of 45-55 s duration were extracted, band-pass filtered (1-15) Hz with a 2^nd^ order Butterworth filter, and delay corrected (∼13 ms). A linear transformation based on this manipulation was obtained per participant[95] to generate spatial filters that correspond to magnetic fields generated by the left and right auditory cortex (S3 Fig). This spatial filter was applied to the raw data and the resulting neural signal, representing the most reproducible component of the evoked data, was selected as a single virtual sensor in analyses henceforth.

### Neural data analysis

#### Spectrotemporal response function of stimulus representation

For multitone patterns, pure onset representations only carry information at a time beginning with the onset of a tone. We formulate this representation as

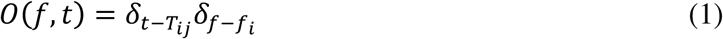

where every onset has equal weight independent of its tone’s frequency band *f*_*i*_ (*i* = 1,…,10), with onset times T_ij_ *T_ij_* of the *j*-th tone with frequency *f_i_; δ_n_* is the discrete unit impulse centered at sample *n*. The input-output relation between this representation of auditory input and the evoked cortical response *r*(*t*) is then modeled by a spectrotemporal response function (STRF). For discrete data this linear model is formulated as:

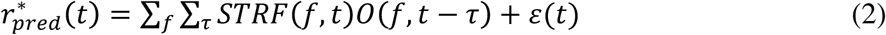

where *ε*(*t*) is the residual contribution to the evoked response not explained by the linear system. Summing only over the frequency term allows evaluating the temporal profile of the response function model (TRF). Exploration of alternative stimulus representations requires substitution of the *O*(*f*,*t*) term in (2) by the analogous time-frequency representation of the stimulus (e.g. by a spectrogram *S*(*f*,*t*)). For all stimuli, *stimulus envelope filterbank* representations were obtained by passing the original waveform through a filterbank of ten order 1000 FIR filters with passbands at mid-values between *f*_*i*_ neighbors (see *Stimuli*, above) starting at 143 Hz. Filter delays were compensated and the envelope in each band was extracted as above. Sampling rates were reduced to 1 KHz and signals smoothed by a delay-corrected 4th order binomial FIR filter. Half-wave rectification (i.e. setting negative values to zero) of the derivative of the stimulus envelope filterbank output gave *envelope onset* representations of the stimulus signal based on the filterbank. Prior to reverse correlation, both envelope and envelope onset representations were transformed to dB-scale.

#### Linear STRF model estimation

STRF estimation was performed via *boosting*, a technique where the error estimate *ε*(*t*) (in Eq. 2) is minimized iteratively via sequential modifications to the STRF[29]. The name originates from the ability to improve (‘boost’) an estimate learning algorithm by establishing aggregate decision rules from across a sequence of many estimation steps, each needing only slightly-better-than-chance accuracy[96–98]. This technique can then be implemented as a forward stage-wise fitting that follows a greedy heuristic, by adding the contribution with the largest available mean-squared-error reduction at each given step[29,99] and in turn maximizing the predictive power of the model[11]. Operationally, STRF estimates by boosting were initialized as a null matrix of dimensions *T*x*F*, where *T* equals the number of experimental time bins and *F* is the total of frequency bins (=10; for TRF estimates, *F*=1); optimization followed through exploring fixed increments and decrements per spectrotemporal bin individually. Among the resulting 2x*F*x*T* possible choices, the outcome with minimum mean-squared-error was selected as the next step in the running STRF estimate. The procedure was iterated, accumulating optimizations, until modifications instead produced a sustained increase in mean-squared error[99], since the method is not guaranteed to find a global optimum. This termination method effectively imposes a sparse structure on the STRF, which allows for extraction of high-temporal resolution features in the STRF even if only low-frequency content was present in the input waveforms (other STRF estimation methods such as normalized reverse correlation[8] and generalized linear models could also used [29,100,101]). Other detailed descriptions of the boosting algorithm implementation for timeseries data, including MEG/EEG are available[29,36].

#### STRF predictive power bounds

The measured evoked cortical response *r*\*(*t*) may include stimulus-independent noise, the presence of which is a consequence of the finite dataset size and leads to STRF model parameters that overfit to the training data. Performance measures that account for stimulus-independent noise are necessarily overestimates and therefore can be considered to act as empirical upper bounds of model performance[11]. In contrast, the risk of overfitting can be minimized using *cross-validation*, where a fraction of the *r*\*(*t*) timeseries r(t) (e.g. 90%) is reserved for model training, and testing is done on the remaining fraction incorporating only the model’s ability to generalize over novel stimulus instances. This would be expected to underperform with respect to an optimal model for the dataset in question and so indicates a lower bound for its performance [11]. In practice, it is this conservative, cross-validated lower-bound that is used for STRF estimates.

#### Nonlinear extension

Linear encoding models may fail to characterize firing rate predictions based on effects such as threshold activity, past-history dependencies, dynamic range compression, synaptic transfer, and the non-negative distribution of the neuron response for example. At the single neuron level, predictions can be improved via introduction of static nonlinearities derived by empirical fit[42], or via intermediate nonlinearities in more complex model hierarchies[101]. For coarse-grained continuous neural responses such as local field potentials and the MEG signal here, it appears that such model hierarchies may no longer apply well. When a static nonlinearity was incorporated using a linear-nonlinear (LN) model[102,103], only a 2% improvement to predictive power resulted (quadratic fit, R^2^=0.972, S5 Fig) and so was not pursued.

#### Estimation of STRF predictive power and noise limit extrapolation

To assess STRF model validity, predictive power was estimated as the fraction of a response signal variance that is stimulus-explained, and corrected for the reduction of noise-related variance achieved by averaging[11]. Namely, for MEG response timeseries *r*_1_(*t*), …, *r_N_*(*t*) r_1_(t), …, r_N_(t) where *N* is the number of repetition trials, total variance is expressed as the average of each trial’s individual variance

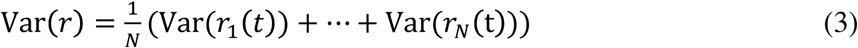

while evoked variance can be expressed as that of the average response Var(*r*). When *N* is large, the extent to which total variance is larger than evoked variance indexes reliability for the response source. Contributions to total variance Var(*r*) are then partitioned into those stemming from the evoked *signal*, and the remainder is treated as *noise*:

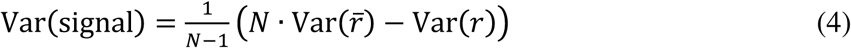

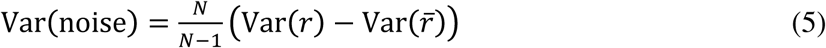

such that estimates are corrected for cases where *N* is small. Often, STRF model estimates are optimized to produce accurate predictions of the evoked response only; in such cases, use of single-trial variance provides an additional statistic regarding the event-related contribution to available recordings. Once a STRF model has been obtained for a particular condition and subject, its ability to predict the evoked response is assessed as the extent of evoked response variance that is not residual error, that is Var(*r*) − Var(*r* − *r*_*pred*_). This expression is the model’s predictive power, which after division by the estimated signal power (eq. 4) Var(signal), represents the fraction of stimulus-evoked variance described by the linear STRF model contingent on a given experimental condition and subject. Analogously, noise power in the same response may be normalized by the estimated signal power, providing the inverse proportion to which the procedure of averaging reduces response variability. When *N* is very large, a normalized noise power of e.g. 10 indicates that averaging reduces variance in the evoked signal to almost a tenth of the original total variance. In the hypothetical case where the procedure of averaging yields no reduction in variability (such as with identical trial response instances), the absence of variability reduction implies an absolute zero noise level. Empirically, each dataset’s (condition and subject) predictive power can be indexed by the intrinsic noise power (e.g. Fig 1C). Assuming the responses have been measured from a similar population, regression analysis may produce an estimate of the STRF model class predictive power, via its extrapolation to the theoretical noise-free limit[11].

## Acknowledgements

We thank Elizabeth Nguyen for excellent technical assistance. This study was funded by the US National Institutes of Health (R01-DC-00843, R01-DC-014085) and the US National Science Foundation (OAC-1063035). We thank support to MVD by the NSF (DGE-142487) and to FCC by the Mexican Consejo Nacional de Ciencia y Tecnología through its graduate scholarship program. The funders had no role in study design, data collection and analysis, decision to publish, or preparation of the manuscript.

